# A simple model explains the cell cycle-dependent assembly of centromeric nucleosomes in holocentric species

**DOI:** 10.1101/2021.04.13.439589

**Authors:** Amanda Souza Câmara, Veit Schubert, Martin Mascher, Andreas Houben

## Abstract

Centromeres are essential for chromosome movement. In independent taxa, species with holocentric chromosomes exist. In contrast to monocentric species, where no obvious dispersion of centromeres occurs during interphase, the organization of holocentromeres differs between condensed and decondensed chromosomes. During interphase, centromeres are dispersed into a large number of CENH3-positive nucleosome clusters in a number of holocentric species. With the onset of chromosome condensation, the centromeric nucleosomes join and form line-like holocentromeres. Using polymer simulations, we propose a mechanism, relying on the interaction between centromeric nucleosomes and Structural Maintenance of Chromosomes (SMC) proteins. All simulations represented a ~20 Mbp-long chromosome, corresponding to ~100,000 nucleosomes. Different sets of molecular dynamic simulations were evaluated by testing four parameters: 1) the concentration of Loop Extruders (LEs) corresponding to SMCs; 2) the distribution and number of centromeric nucleosomes; 3) the effect of centromeric nucleosomes on interacting LEs; and 4) the assembly of kinetochores bound to centromeric nucleosomes. We observed the formation of a line-like holocentromere, due to the aggregation of the centromeric nucleosomes when the chromosome was compacted into loops. A groove-like holocentromere structure formed after a kinetochore complex was simulated along the centromeric line. Similar mechanisms may also organize a monocentric chromosome constriction, and its regulation may cause different centromere types during evolution.

## INTRODUCTION

Centromeres are required for the movement of chromosomes during cell division. Most organisms contain a single size-restricted centromere per chromosome (monocentromere), visible as a primary constriction during metaphase. However, in independent eukaryotic taxa, including some protists, plants, and invertebrates, species with holocentric chromosomes have evolved repeatedly (1–3).

Holocentric chromosomes have no distinct primary constriction visible at metaphase. Instead, spindle fibres are attached along almost the entire poleward surface of the chromatids (4). Due to the chromosome-wide distribution of holocentromeres, single-chromatids cohere along the entire chromatids and appear as two parallel structures at metaphase. In contrast, in monocentric species, the sister chromatid cohesion is restricted to a single position at the centromere and X-shaped metaphase chromosomes are formed.

Clades possessing holocentromere types include more than 350,000 species in total (5). Likely, holocentricity is even more common than reported so far as the identification of the centromere type is challenging for small chromosomes. One common explanation for the development of holocentric chromosomes during evolution is their putative advantage to tolerate DNA double-strand breakages inducing chromosomal fragments, which will not be lost during cell division (6).

Different types of holocentromeres exist, as exemplified by either presence or absence of the centromere-specific histone H3 variant CENH3 (also called CENPA) and centromere-specific repeats, different morphology of chromosomes and diversity of meiotic behaviour (4, 7, 8). The mechanism that gives rise to holocentricity is still unknown. However, the fact that holocentrics arose independently several times during evolution suggests that the transition from mono- to a holocentromere type may be a relatively simple process.

In contrast to most monocentric species, where no obvious dispersion of the centromeres occurs during interphase, the organization of holocentromeres differs between interphase and mitotic metaphase (Figure 1). During interphase, e.g. in the nematode *Caenorhabditis elegans* (9) and plant species, the Juncaceae *Luzula elegans* (10, 11) and the Cyperaceae *Rhynchospora pubera* (12) holocentromeres are dispersed into a large number of CENH3-positive centromeric nucleosome clusters, which are evenly distributed within the nucleus. With the onset of chromosome condensation, the centromeric nucleosome clusters join and form line-like structures along both chromatids. After segregation of chromatids, dispersion of holocentromeres is concomitant with chromatin decondensation. Hence, the organization of holocentromeres is cell cyle-dependent and the line-like metaphase centromere is a result of fused centromeric nucleosomes. Due to this multi-centromere subunit structure, holocentric chromosomes could also be considered as ‘polycentric’ as proposed by (13). However, also a monocentromere is composed of multiple centromeric nucleosomes based on the centromere subunit model, where the centromere is assembled from repetitive subunits tandemly arranged on a continuous chromatin fibre (14). An additional centromere feature of the holocentric genera *Luzula* and *Rhynchospora* is the presence of a longitudinal groove along the poleward surface of mitotic metaphase chromosomes (Figure 1) (10–12, 15). However, this centromere structure does not exist in all holocentric species (12, 16).

**Figure 1.**
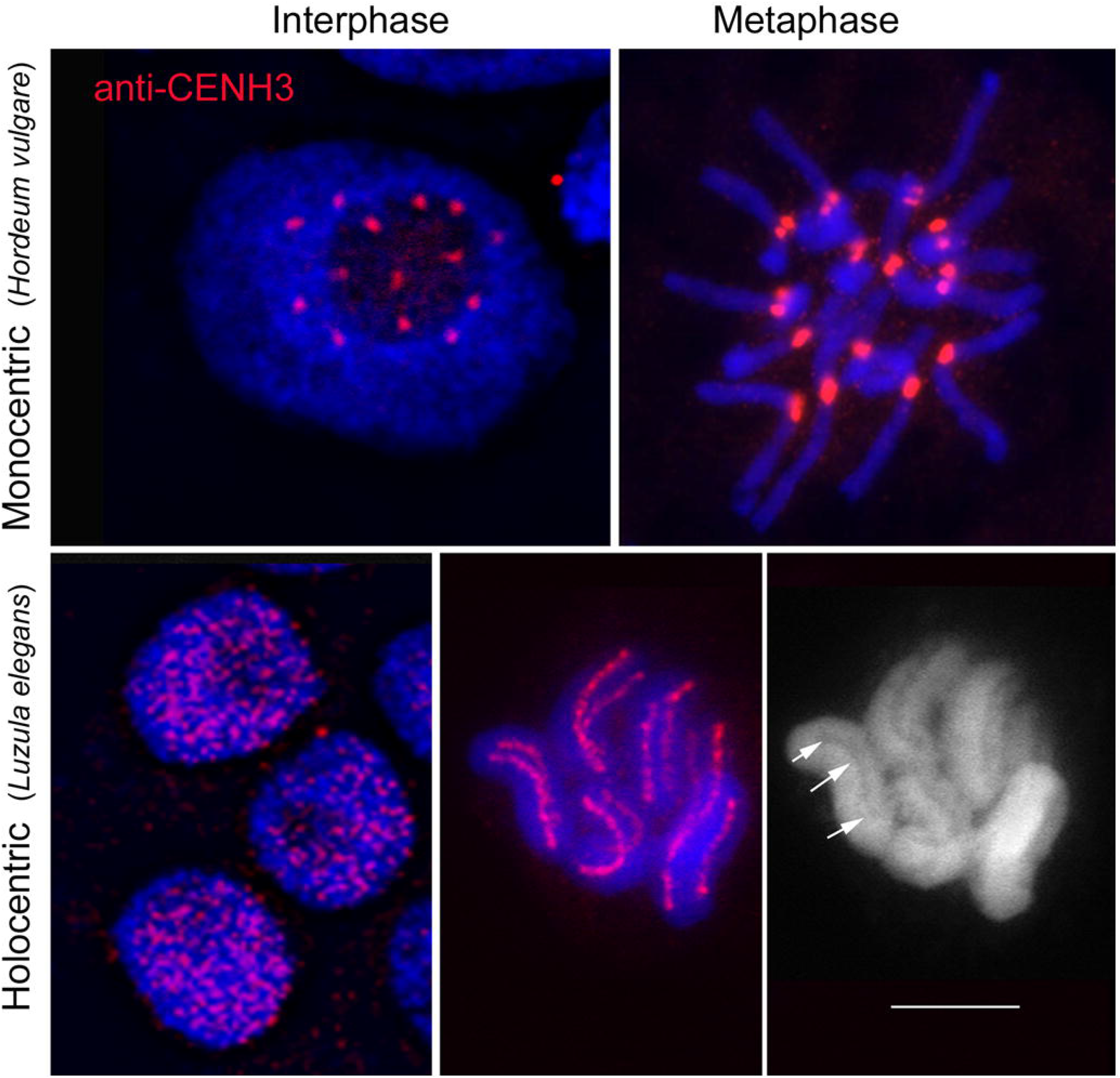
Interphase and metaphase chromosomes of monocentric (*Hordeum vulgare*) and holocentric (*Luzula elegans*) species. During interphase, e.g. in *L. elegans*, holocentromeres disperse into a large number of CENH3-positive centromeric nucleosome clusters. With the onset of chromosome condensation, the centromeric nucleosome clusters join and form line-like structures along both chromatids. In contrast, in most monocentric species, e. g. in *H. vulgare* no obvious dispersion of the centromeres occurs during interphase. CENH3 (red) indicates the position of centromeres. Arrows indicate the longitudinal centromere groove in *L. elegans*. Chromosomes were counterstained with DAPI. Bar = 5 μm. Pictures were taken from Heckmann et al. (11) with permission from S. Karger AG, Basel.

The peculiar cell cycle dynamics and structure of holocentromeres prompt us to apply a loop extrusion model to decipher the potential mechanism behind the cell cycle-dependent assembly of centromeric nucleosomes and the formation of a groove-like centromere structure. It has been shown that the extrusion of chromatin loops affects the condensation and segregation of sister chromatids (17). Loop extrusion relies on the action of Structural Maintenance of Chromosomes (SMC) complexes. SMC proteins are a group of evolutionarily conserved protein complexes including cohesins, condensins and SMC5/6 complexes sharing similar structures and dynamics (18, 19). They bind to the DNA molecule and translocate along with it, progressively bringing together loci separated by larger distances in the chromosome and leaving a DNA loop behind (20–22). But neither do we know the exact mode of action of SMC complexes nor is it clear whether differences exist between species (18).

The motion of SMC complexes can be affected by other proteins such as the CCCTC-binding factor (CTCF) or the wings apart-like protein (WAPL), which were reported to anchor or release the approaching SMC complexes (18). By anchoring, for example, CTCFs are proposed to fix loop bases, thus delimiting regions highly self-contacting, called Topological Associated Domains (TADs) (23). SMC complexes themselves can also block each other, leading to the formation of more compact loop arrangements (24). Thus, varying SMC complex concentrations can generate different chromatin condensation levels.

In mitotic chromosomes, cohesins are related to sister chromatid cohesion, while the condensation of chromosomes relies on the function of condensin I and II (25). In invertebrates, the two condensins have been associated with different phases of the cell cycle. Condensin I becomes active after nuclear membrane breakdown, whereas condensin II is active already in G2 (26).

In monocentric species, condensin II acts on the axial shortening of the chromosome, while condensin I acts on the lateral compaction (25). Originally, condensin I was believed to be lost and not required in holocentric species (27) until its later discovery in *C. elegans* (28). Nonetheless, the condensation of mitotic chromosomes is mainly attributed to condensin II, whose depletion profoundly affects prophase condensation. Condensin II is present along the holocentromeres of mitotic *C. elegans* chromosomes, but condensin I appears dispersed, and its depletion did not cause prophase condensation defects (28). The occurrence of SMC complexes is possibly a general feature of holocentric chromosomes, as the cohesion α-kleisin subunit also colocalizes with the holocentromere of the plant *L. elegans* (29).

In this work, we simulated the cell cycle-dependent formation of a holocentromere-like chromosome based on the condensation of a single chromatin fibre possessing a large number of centromeric nucleosomes. To keep our simulation as simple as possible, we considered only the compaction by a general SMC complex type, what we called Loop Extruder (LE). Additionally, we applied possible chromatin fibre crossings of the chromatin fibre, mimicking the action of topoisomerase II, as described in (24). Other factors involved in the process of chromosome condensation were ignored. The LE worked as postulated in the loop extrusion model of (30). When the extruded loops grow larger, more distant DNA regions can interact, allowing chromatin domains to be compacted by loops. We assumed a scattered distribution of centromeric nucleosomes, proposing they affect the LEs motion. Like CTCF and WAPL proteins, centromeric nucleosomes were already suggested to affect the loop extrusion process because the compaction of chromatin is interrupted at centromeres with chromatin loops limited to the pericentromeric region (31, 32).

We performed different sets of molecular dynamic simulations by testing four parameters: 1) the concentration of LEs; 2) the distribution and number of centromeric nucleosomes; 3) the effect of centromeric nucleosomes on interacting LEs; and 4) the assembly of kinetochores bound to centromeric nucleosomes. We observed the formation of a line-like holocentromere, due to the aggregation of the centromeric nucleosomes when the chromosome was compacted into loops. A groove-like holocentromere structure was formed after a kinetochore complex was simulated along the centromeric line.

## MATERIALS AND METHODS

All simulations represent single ~20 Mb-long chromosomes, modelled as a polymer chain with 100,000 monomers. Each monomer corresponded to one nucleosome. Centromeric nucleosomes are uniformly distributed along the chromosome. They differed from noncentromeric nucleosomes only regarding the interaction with the simulated Loop Extruders (LEs). The LE was simulated as a dynamic bond between two nucleosomes.

### One-dimensional (1D) simulations of loop extrusion

We performed 1D simulations to determine which nucleosomes the LEs bind as a function of time. ln the simulation, the LEs initially bound pairs of adjacent nucleosomes. There was always the same number of LEs during the simulation. The nucleosome pairs that LEs bind changed according to the extrusion motion and the interaction rules.

For the extrusion motion, we applied the algorithm of Alipour and Marko (30) and Goloborodko et at. (24). We adopted two-sided LEs because (33) showed that one-sided LEs could not reproduce alone some biological phenomena. As a two-sided extrusion motion, both nucleosomes bound by a LE progressively changed. The left-side nucleosome always changed to the one on its left and the right-side nucleosome to the one on its right. This change occurred once every 1D step, which is the velocity of the LE. With this extrusion motion, the LE bound progressively more distant nucleosomes, until the LE unbound and reinitiated its motion at another side, with a new pair of nucleosomes. This occurred with a chance of one over 1,000 1D steps, a period called lifetime.

Two interaction rules affect the extrusion. One nucleosome cannot be bound by more than one LE, and adjacent LEs block each other’s way, but can continue the extrusion in the opposite direction, i.e. that not occupied by another LE. When a LE meets a centromeric nucleosome one of the three settings apply: 1) no effect – the LE continues its motion freely; 2) blocking effect – the LE motion is blocked on one side; and 3) anchoring effect – the LE is blocked and not allowed to unbind, even past its lifetime. In all settings, the centromeric nucleosomes were not allowed to be occupied by a LE.

The final list of nucleosomes bound by LEs overtime was later used to (i) calculate the chromosome and average chromatin loop lengths; (ii) verify if the simulation has reached a compaction equilibrium; and (iii) pass the data over to three-dimensional simulations. We calculated the chromosome length as the total number of nucleosomes outside loops, where the nucleosomes bound by a LE counted as outside the loop. For example, if there were no chromatin loops, then the chromosome length was simply the total number of nucleosomes along the chromatin fibre. If there was a loop binding two distant nucleosomes, then the minimum distance was shortened by the chromatin loop length. The average chromatin loop length was calculated as the sum of nucleosomes between each pair of LE-bound nucleosomes over the number of LEs. We considered that the simulation reached an equilibrium when these two parameters were stable over time. To verify this, we performed 10 replicates of each model.

### Three-dimensional (3D) polymer simulations of single chromosomes

We performed Langevin dynamics simulations with OpenMM Python API (Application Programming Interface) (34), using the integrator VariableLangevinIntegrator with 300 K temperature, 0.001 ps^−1^ friction coefficient and 80 fs error tolerance. We used the python library from https://github.com/mirnylab/ to implement simulation parameters. The following three forces composed the force field of the chromosome: 1) harmonic force between adjacent nucleosomes, with 10 nm mean distance between them and 1 nm wiggle distance – this conferred the chromatin fibre of 10 nm thickness; 2) Grosberg stiffness force with 1 k_B_T stiffness constant; and 3) repulsive polynomial force up to 10.5 nm distance, that allowed crossing of the chromatin fiber over 5 k_B_T energy – mimicking the presence of topoisomerase II, as in (35). This force field was valid to all centromeric and non-centromeric nucleosomes.

The list of nucleosomes bound by LEs over time (retrieved from the 1D simulation) integrated the loop extrusion into the 3D simulations. The 1D simulation accounted for the effects of the centromeric nucleosomes in the loop extrusion, and this was enough to distinguish them from the other nucleosomes. The binding of nucleosomes by LE was simulated as a harmonic force with 5 nm mean distance and 0.5 kBT/nm^2^ harmonic force constant, as in (32). This force was updated every block of 3D steps (100 3D steps), according to the bonds list from the 1D simulation, where 1D step corresponds to a block of 3D steps. The simulation started with a random conformation, run the first 10,000 blocks without loop extrusion and then run 40,000 more blocks with a constant number of LEs.

### 3D simulations with a kinetochore

This simulation presented two different objects to which different forces were applied, chromatin and the kinetochore. Chromatin was simulated as above. The kinetochore was simulated by fixing 25,000 non-connected beads on a regular grid of 50 × 250 × 2000 nm^3^. Only two forces acted upon the kinetochore beads: a tethering harmonic force with 5 k_B_T/nm^2^ spring constant; and a polynomial repulsive force as before, but with 10,000 k_B_T energy truncation – so the chromatin fibre could not cross the kinetochore. Centromeric nucleosomes were also tethered during the simulation. In the initial conformation, the centromeric nucleosomes were aligned along the z-axis, and non centromeric nucleosomes formed straight chromatin loops along the x-axis. The data for these loops were taken from 1D simulations of loop extrusion. The 1D simulation run for 100,000 1D steps, but only the last 50,000 were used in the 3D simulations with the kinetochore. This allowed the system to equilibrate while still performing loop extrusion.

## RESULTS

### Prerequisites of the model

We simulated the cell cycle-dependent condensation process of a modelled holocentric chromosome to test factors involved in the line-like assembly of centromeric nucleosomes during mitosis. All simulations represented a ~20 Mbp-long chromosome, corresponding to ~100,000 nucleosomes. Chromosomes of this length exist in the holocentric worm *C. elegans* (36).

Our dynamic model was based on the following assumptions. The chromosomal 10 nm chromatin fibre is represented as a beads-on-a-string polymer (Figure 2a), in which each bead corresponds to one nucleosome containing two copies of histone H3, H4, H2A and H2B. We assumed that ~ 200 bps of DNA represent 147 bps wrapped around each nucleosome plus the nucleosome linker DNA (24). Ring-like SMCs were simulated as chromatin fiber loop extruders (LEs). They progressively bind distant nucleosomes but can also unbind from the chromatin fiber (Figure 2b). When two LEs meet during the chromatin loop extrusion process, they block each other’s way, as proposed by (24, 30), thus generating side-by-side and nested loop arrangements, as shown in Figure 2c. Dynamic binding and release of LEs from the chromatin fibre resulted in a compacted mitotic chromosome, in agreement with the simulations of Goloborodko et al. (24).

**Figure 2.**
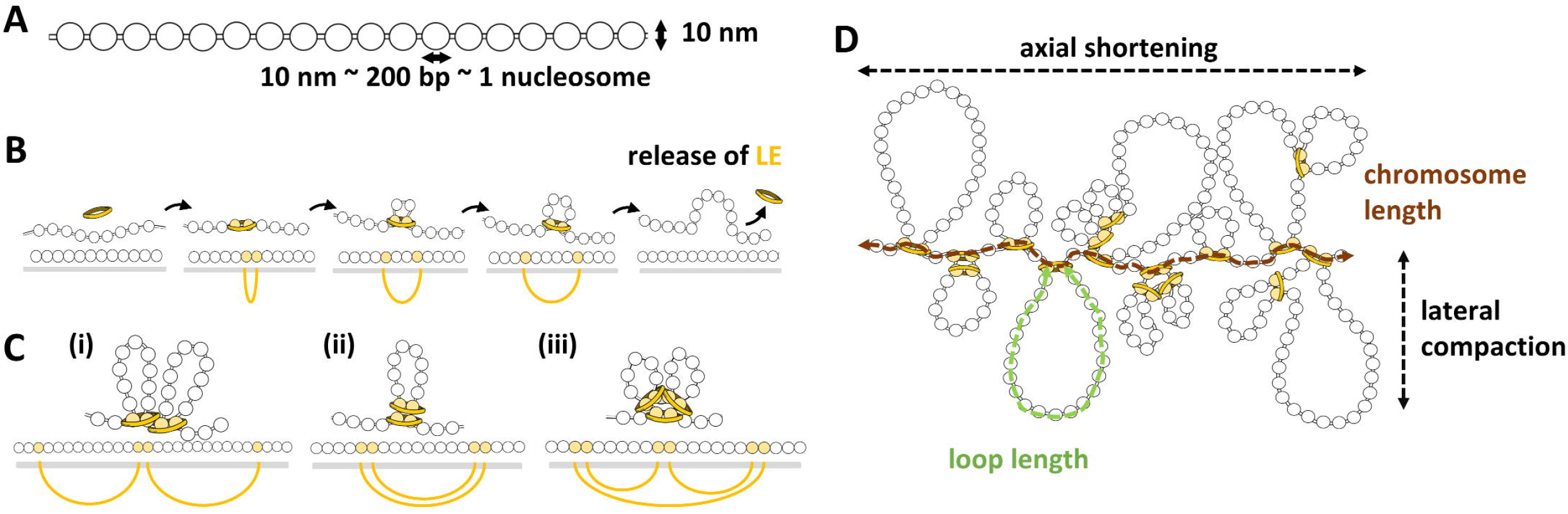
Schematic representation of the adopted chromosome and loop extrusion models. A) The chromosomal 10 nm chromatin fibre represented as a beads-on-a-string polymer. Around ~200 base pairs of DNA (including the linker) are wrapped around each nucleosome. B) The loop extrusion model. The Loop Extruder (LE) is represented by a yellow ring. Nucleosomes bound by LEs are shown in yellow, the bond between them is represented as a yellow ellipsoid line, and the grey bar represents the chromatin. C) Different examples of loops formed by two proximal LEs; (i) side-by-side loops, (ii) nested loops and (iii) combination of both. D) Chromosome condensation by loop formation. The bases of the loops form the axis of the chromosome, and the loops are radially distributed. The degree of chromosome condensation is due to axial shortening and lateral compaction. These two parameters are functions of the nucleosome number and can be computed as the chromosome length and the average loop length (measured as the number of nucleosomes), respectively.

The degree of chromosome condensation can be determined by two parameters - lateral compaction and axial shortening of chromosomes (25). We associated these two parameters to the average chromatin loop and chromosome lengths, respectively (Figure 2d). We defined the chromosome length as the total number of nucleosomes outside the loops and the loop length as the nucleosomes inside the loop, regardless of centromeric or non-centromeric nucleosome. These two parameters reached an equilibrium in the later stages of the simulation (Supplementary Figure 1), representing a condensed chromosome. We used both parameters to compare quantitatively the degree of chromosome condensation of final conformations obtained after different simulations.

The dynamic behaviour of centromeric nucleosomes is an integral part of our model. We randomly chose the positions of centromeric nucleosomes, uniformly distributed, to mimic holocentric or monocentric chromosomes. The only unique feature of centromeric nucleosomes was their ability to interfere with the motion of LEs.

The release of a LE was determined by its lifetime, a computational parameter that relates to the experimental affinity of condensins to the DNA. But the interaction with centromeric nucleosomes might alter the dynamics of the LEs (31). Other proteins were already found to interfere with the motion of SMCs. CTCFs, for example, anchor cohesins and prevent their release (37). WAPL releases cohesin and prevents the extension of chromatin loops (38).

We considered two different interaction effects of the centromeric nucleosomes (Figure 3) and compared them to a lack of effect, which allowed LEs to pass freely, as at non-centromeric nucleosomes. As a barrier of LEs, a centromeric nucleosome partially blocks the motion of the LE. As shown in Figure 3A, when a LE meets a centromeric nucleosome, the motion of the LE in the direction towards the centromeric nucleosome is blocked. In the opposite direction, the LE continues to reel and to extrude a loop until the eventual LE release. As an anchor of LEs, a centromeric nucleosome partially blocks the motion and prevents the release of the LE. When a LE meets a centromeric nucleosome, it continues to reel in the opposite direction until meeting another LE or centromeric nucleosome (Figure 3A). For comparison, Figure 3B exemplifies a loop extrusion by one LE if centromeric nucleosomes would have no effect in its motion.

**Figure 3.**
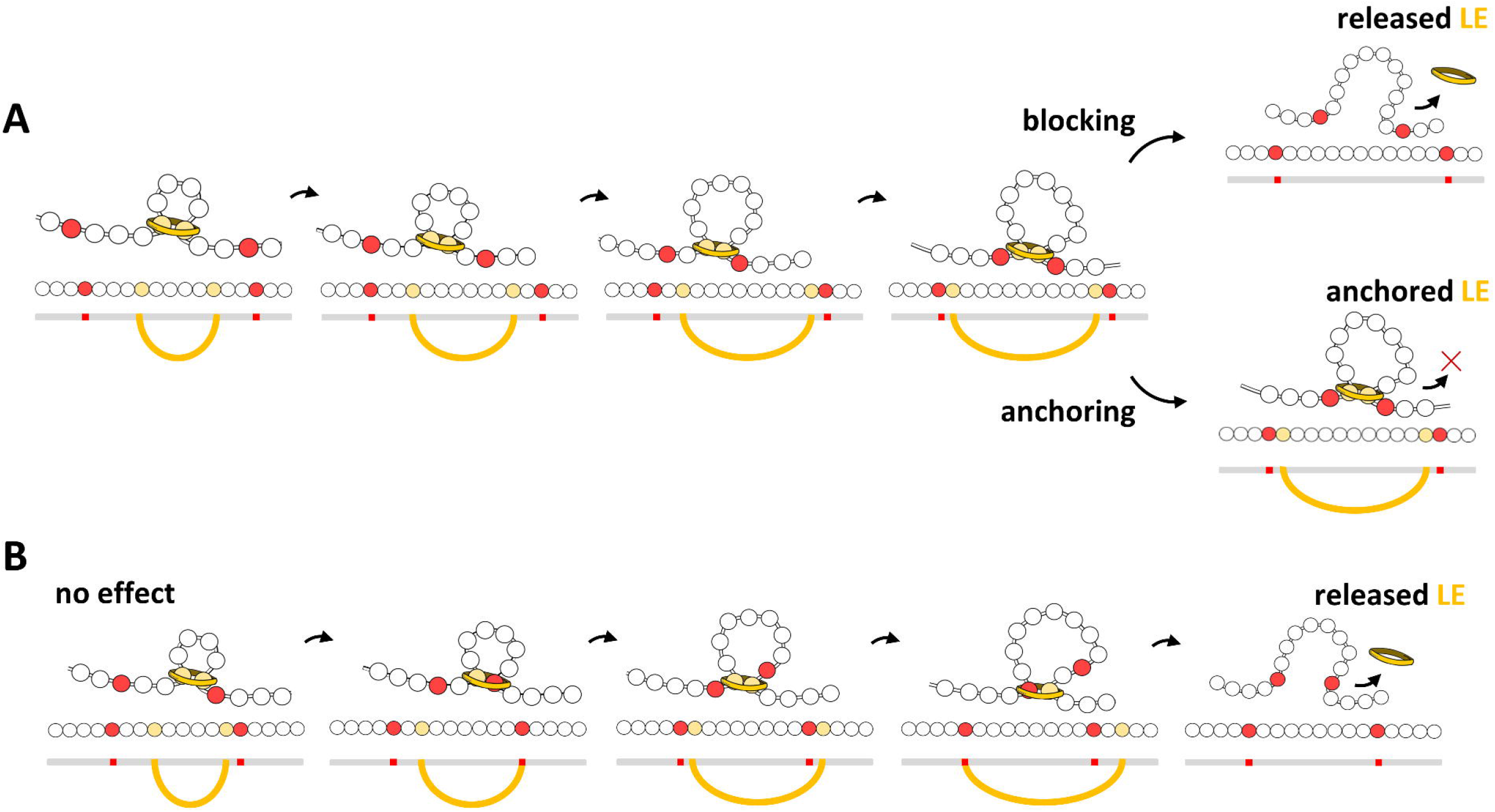
Effects of the presence of centromeric nucleosomes (in red) in the loop extrusion process. A) Blocking and anchoring effects. In both, one centromeric nucleosome blocks the motion in its direction, but the opposite side of the LE (in yellow) continues to reel. In the blocking effect the LE interacting with the centromeric nucleosome can unbind, but in the anchoring effect the LE is permanently bound to the centromeric nucleosome. B) For comparison, centromeric nucleosomes that do not interact with the LE can pass through it.

Besides, we considered the cell‐cycle dependent, centromeric localization of kinetochore proteins. The kinetochore is a multiprotein complex that connects centromeric nucleosomes to the microtubules and is considerably larger than a nucleosome. The composition of the kinetochore is cell cycle-dependent and forms at both metaphase chromatids, a plate-like structure at the side of centromeric nucleosome clusters (1, 39, 40). Thus, we simulated an orderly arrangement of kinetochore units, as a set of beads with fixed positions in a grid forming a chromosome-wide, plate-like structure. Then, we tethered the centromeric nucleosomes to one side of this kinetochore arrangement, leaving the opposite side free to interact with microtubules, although microtubules were not simulated. Table 1 lists the simulation parameters we varied in this work, and the three hypotheses on the arrangement of centromeric nucleosomes.

**Table 1.** Main features of the proposed hypotheses and tested parameters

| **Hypothesis on** | **Number of**<br>**LEs** | **Centromere type** | **Interaction effects**<br>**of centromeric**<br>**nucleosomes** | **Kinetochores**<br>**structure** |
| --- | --- | --- | --- | --- |
| ***Formation of the***<br>***centromeric line*** | 50 - 2000 | Holocentromere | None, blocking, or<br>anchoring | Not considered |
| ***Different centromere***<br>***types*** | 1000 | Mono- and<br>holocentromere | Anchoring | Not considered |
| ***Holocentromeric***<br>***groove*** | 1000 | Holocentromere | Anchoring | Plate-like grid of<br>beads |

### Anchoring of Loop Extruders (LEs) by dispersed centromeric nucleosomes leads to the formation of a line-like holocentromere during chromosome condensation

We tested sixteen different amounts of LEs (between 50 and 2000) interacting with a single 20 Mb-long chromatin fiber, containing 100 uniformly distributed centromeric nucleosomes. A comparable distribution exists in *C. elegans* (41). We quantified the degree of chromosome condensation relative to the amount of LEs by calculating the average chromatin loop and chromosome lengths (Figure 4a).

**Figure 4.**
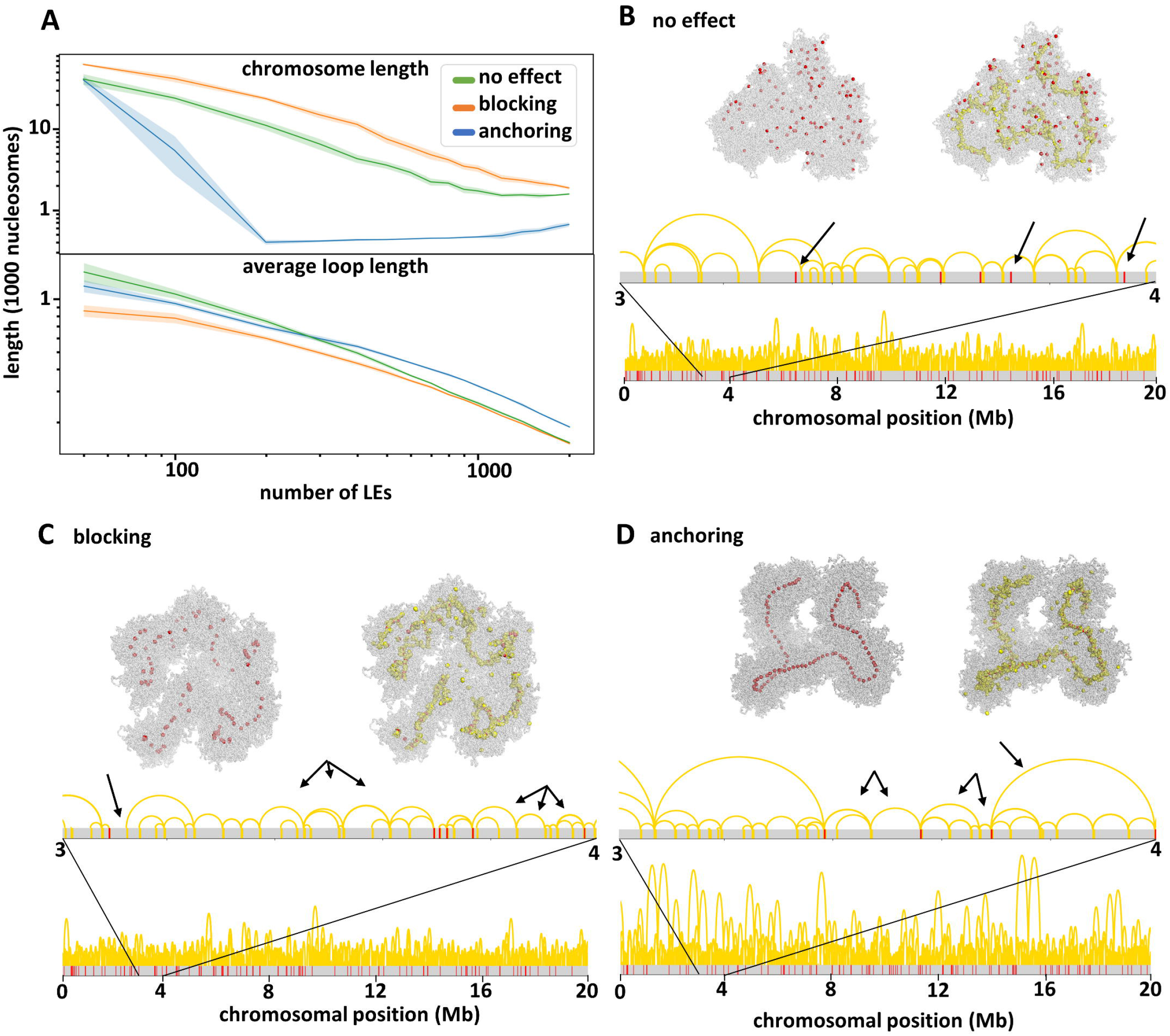
Effects of centromeric units. A) Chromosome length and average loop length as a function of the number of simulated LEs. Both parameters were calculated from simulations considering three different effects (no effect, blocking or anchoring) of centromeric units in the loop extrusion process. (B – D) Final conformations of three simulations with different centromeric effects (see Supplementary Movies 1-3 for simulation examples). The distribution of centromeric nucleosomes (red) and LEs (yellow) is shown in the chromatin fibre (grey), in the 3D structure (top) and sequence (bottom). For each conformation arrows indicate characteristic loop organizations. (B) With no effect, loops are observed spanning centromeric nucleosomes. (C) With the blocking effect, regions are observed outside loops as well as multiple loops between two adjacent centromeric nucleosomes. (D) With the anchoring effect, only one or two loops are observed between adjacent centromeric nucleosomes.

We first considered the setting where centromeric nucleosomes made no effect on the loop extrusion process (Supplementary Movie 1). We observed that the presence of LEs alone could bring the chromosome to a condensed state, as reported in (24, 30). The more LEs were considered, the more condensed was the chromosome, and both chromosome length and average loop length decreased. But after compaction, centromeric nucleosomes were dispersed and not arranged in a holocentromere-like manner. LEs concentrated in the axis of the chromosome, forming the basis of side-by-side and nested loops that span centromeric nucleosomes (Figure 4b).

Then, we considered that centromeric nucleosomes blocked the motion of LEs (Supplementary Movie 2). Again, an increase in the number of LEs consistently decreased the average chromatin loop length, bringing the chromosome into a more condensed state (Figure 4c). Centromeric nucleosomes were arranged in small groups, preferentially central to the chromosome axis. These centromeric nucleosomes were brought together by the blocking effect of the LEs, becoming a focal point of LE accumulation. This grouping of centromeric nucleosomes was only transient, since LEs were able to become released. In contrast to the model where centromeric nucleosomes had no effect on LE, we always observed the centromeric nucleosomes outside or at the boundaries of chromatin loops.

Last, we considered that centromeric nucleosomes anchored LEs, i.e. prevented them from unbinding from the chromatin (Supplementary Movie 3). The length of the chromosome and chromatin loops again decreased with the increase of the LE number. But with 200 LEs, the chromosome length reached a plateau of ~500 nucleosomes (Figure 4a). This number corresponds to 100 centromeric nucleosomes arranged in between of 400 LE-bound nucleosomes, which were the basis of 200 chromatin loops. These loops were anchored by LEs and organized side-by-side (Figure 4d, arrowed). Due to the persistent LE anchoring, the proximity of the centromeric nucleosomes became permanent instead of transient. At the end of the condensation process, the entire chromosome was folded into chromatin loops. Inbetween two centromeric nucleosomes, there were always one or two chromatin loops. With the anchoring effect of centromeric nucleosomes, it was possible to model a holocentromere-like structure formed by side-by-side centromeric nucleosomes. In the simulations considering the anchoring effect, the chromosome length remained approximately the same with 200 to 1200 LEs. 200 LEs were enough to anchor all (100) centromeric nucleosomes into a line. The remaining LEs accumulated around the holocentromere-like line and further divided the anchored chromatin loops into smaller loops, leading to a lateral compaction of the chromosome. More than 1200 LEs slightly increased the chromosome length, and the centromeric nucleosomes occurred more distantly to each other. This indicates that many LEs may hinder the formation of a holocentromere-like organization by the accumulation of LEs between centromeric nucleosomes.

When a holocentromere-like structure was formed, the centromeric nucleosomes were linearly organized, meaning that their position along the line followed their position along the chromatin fibre (Figure 5). The chromatin between centromeric nucleosomes was arranged into loops, whose bases were held together by LEs. Chromatin loops were also linearly organized, as expected for mitotic chromosomes (42). Compared to the other settings, where the centromeric nucleosomes had no effect or only blocked the loop extrusion motion, the anchoring of LEs by centromeric nucleosomes resulted in a more than two-fold reduction of the chromosome length, but with similar chromatin loop lengths (Figure 4a). We conclude that the centromeric nucleosomes modulate the length of the condensed chromosome. This result is consistent with the report by Maddox et al. (43), who observed an unusual condensation of chromosomes by depleting centromeric nucleosomes in *C. elegans.*

**Figure 5.**
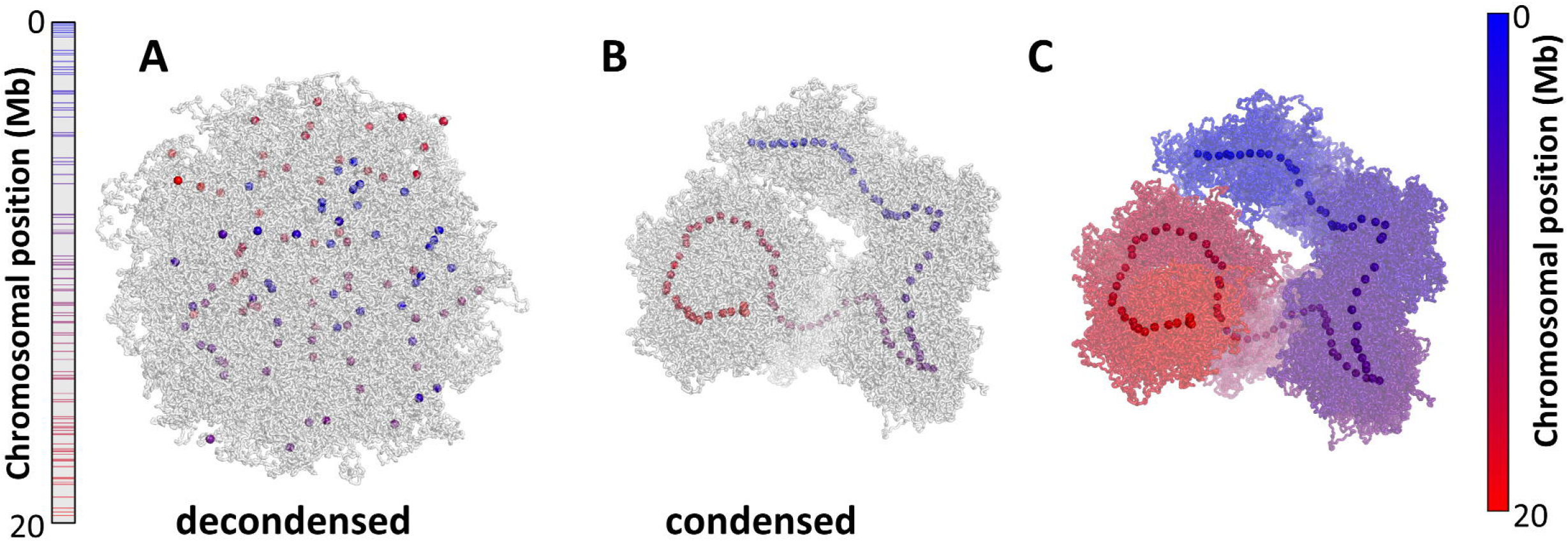
Simulated conformations of a holocentric-like chromosome with 100 centromeric nucleosomes with anchoring effect after condensation by 1000 LEs (see Supplementary Movie 3 for the entire simulation). Centromeric nucleosomes are colored from blue to red according to the position in the linear genome. Left bar represents the chromosome length and colored lines indicate the position of centromeric nucleosomes. (A) The decondensed conformation represents an interphase with dispersed centromeric nucleosomes. (B) Condensation of a chromosome due to loop extrusion. Centromeric nucleosomes are aligned in the axis of the chromosome following the position in the chromosomal sequence. (C) Condensed holocentric chromosome colored from blue to red, as indicated by the bar at the right. The chromosome is entirely linearly arranged along the chromosome axis, following the position in the chromosomal sequence.

### Clustered distribution of centromeric nucleosomes induces a monocentromere-like structure

We modelled the structure of holo- and monocentric chromosomes by changing the distribution and number of centromeric nucleosomes (Supplementary Movies 3 and 4). In the holocentric, 100 centromeric nucleosomes were normally distributed over the entire chromosome; and in the monocentric, 20 centromeric nucleosomes were clustered in a small region corresponding to 400 kb. In both cases, we simulated a ~20 Mb long chromosome containing 1000 LEs.

Non-compacted (interphase-like) and compacted (prophase-like) chromosome conformations for both centromere types were simulated (Figure 6). We only considered the prophase stage, because highly compacted mitotic chromosomes arise from more than one condensation step (35, 43). The non-compacted holocentric chromosome presented abundant and scattered centromeric nucleosomes. In contrast, the centromeric nucleosomes of a monocentric chromosome were clustered. The compacted holocentric chromosome displayed a line of centromeric nucleosomes, whereas the monocentric chromosome had a centromeric region with distinct compaction. The exact arrangement of the chromatin fibre in the centromeric constriction is unknown but loops in our model were smaller in this region (Figure 6, Supplementary Figure 2). The smaller loops arose from the short distance between centromeric nucleosomes, which restricted the extrusion of loops. Compaction of regions without centromeric nucleosomes results in larger chromatin loops (Figure 6), and the differences in loop size created a constriction-like structure.

**Figure 6.**
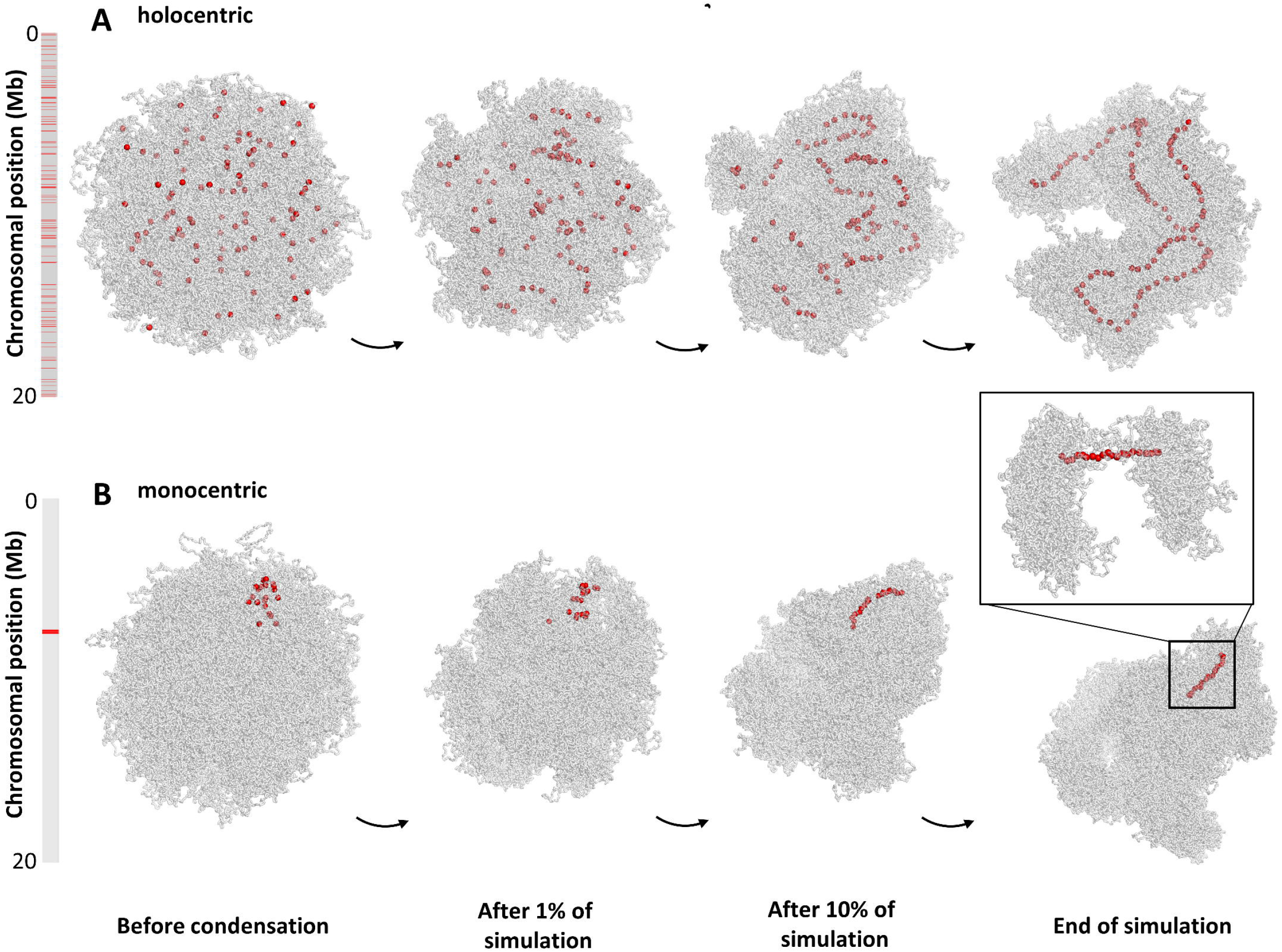
Comparison between simulated (A) holocentric- and (B) monocentric-like chromosomes at different stages of the condensation process (Supplementary Videos 3 and 4). Bars indicate chromosome length and red lines the position of the centromeric nucleosomes. (A) The condensed holocentric-like chromosome presents an average loop size of 325 nucleosomes and a chromosome length of 507 nucleosomes. (B) The condensed monocentric-like chromosome presents loop sizes of 59 and 260 nucleosomes inside and outside the centromeric region, respectively, and a chromosome length of 2230 nucleosomes. The inset shows the centromeric region, with smaller chromatin loops, resembling the centromere constriction of monocentric chromosomes.

Holocentric and monocentric-like compacted chromosomes differed also in length. In line with our previous tests for centromeric nucleosomes with different interaction effects (Figure 4), the monocentric chromosome was about three times longer than the holocentric chromosome (1,760 and 470 nucleosomes, respectively, Figure 6). The monocentric chromosome, with a lower number of centromeric nucleosomes, resembled the setting of no effect when the chromosome length was mostly limited by the number of LEs. The holocentric chromosome, with 100 centromeric nucleosomes, resembled the setting of the anchoring effect when the chromosome length was mostly limited by the number of centromeric nucleosomes.

### The presence of a kinetochore complex might create a mitotic groove-like centromere structure in holocentrics

A longitudinal groove along each miotic sister chromatid is visible in holocentric chromosomes of *L. elegans* and *R. pubera* (10–12). We speculated that the kinetochore, composed of protein layers, restricts the LEs in space, giving a preferential direction to the emergence of chromatin loops. To test this hypothesis, we simulated the kinetochore as a large regular grid of beads next and parallel to the centromeric nucleosomes, which were constricted to a straight line.

We observed in simulations that LEs, and the emergence of loops in the three-dimensional space were restricted by the kinetochore. The chromatin fiber, organized in loops, was free to diffuse but was not able to occupy the entire space opposite to the kinetochore arrangement, thus forming a groove-like structure, and maintained the centromeric line of the chromosome (Figure 7). At the bottom of the groove, we observed associated and aligned centromeric nucleosomes and the surrounding LEs (Figure 7B, C). A transversal cut confirmed the groove-like structure (Figure 7C). Loops surrounding the longitudinal centromere created a clear contrast in the structure and covered the ends of chromosomes.

**Figure 7.**
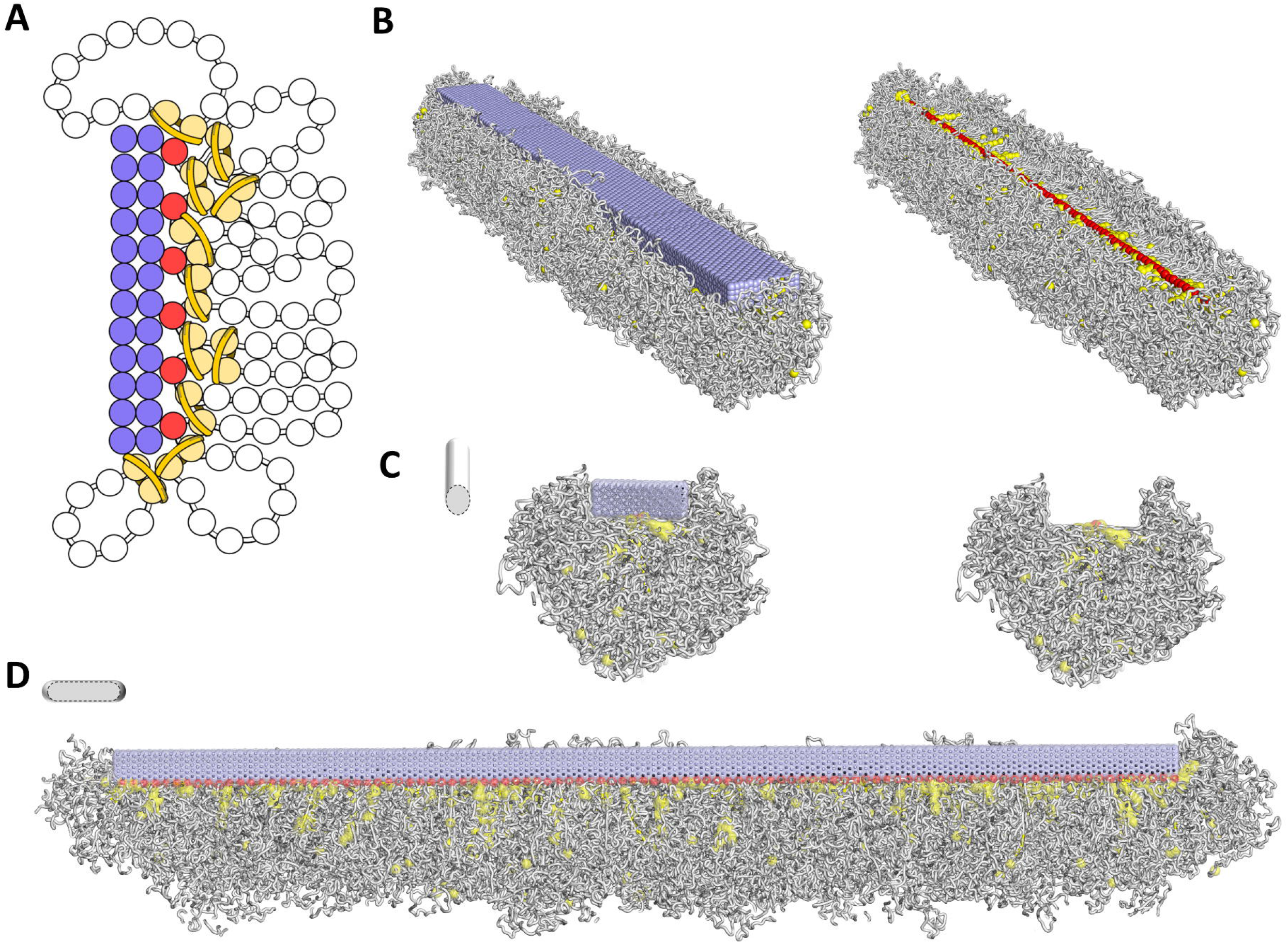
Simulation of a groove-like centromere along a holocentric condensed chromosome. A) Representation of a simulated kinetochore arrangement as a rectangular bar formed by fixed spheres (in lilac). Red spheres represent the line of centromeric nucleosomes. Pairs of yellow spheres represent nucleosomes bound by LEs and the chromatin fibre is shown as white beads on a string. B) final conformation of a simulated holocentric chromosome in the presence of the kinetochore arrangement. Components follow the same code color as in (A). The kinetochore is embedded in the chromatin fibre. On the right the kinetochore is not shown so that the centromeric line is visible at the bottom of the groove and surrounded by LEs. Lateral (C) and longitudinal (D) cross-sections evidence the centromeric groove-like structure.

## DISCUSSION

We propose a loop extrusion process, in which centromeric nucleosomes block and anchor LEs and are brought together into a line in compacted holocentric chromosomes. This mechanistic model relies on the function and distribution of three protein complexes, broadly observed across eukaryotes: SMC proteins, CENH3-containing nucleosomes and kinetochores, which can be regulated to create variable centromeric arrangements.

SMCs, such as condensins, have been shown to act as loop extruders in the chromatin compaction process (44). Their dynamic binding and release of the chromatin fibre induce a stable condensed state of the chromosome (24). The motion of condensins can be affected by other proteins to approximate specific DNA sites and to form regions with distinct condensation, such as TADs containing CTCFs (45, 46). Chromatin loops with restricted size also characterize heterochromatin and pericentromeric regions in mitotic chromosomes (32, 47). Our simulations showed that the alignment of the centromeric nucleosomes occurred only when they were anchored to LEs (Figure 4). The anchor between centromeric nucleosomes and condensins relates to the affinity between them, so we propose that they must present a high affinity. SchaIbetter et al. already suggested that centromeric nucleosomes could act as barriers to the loop extrusion motion (31). The affinity between condensins and chromatin was also reported (48), and it is possible that in complexes with centromeric nucleosome, this affinity is even higher.

In our model, similar to TAD borders, adjacent centromeric nucleosomes had a high contact probability in the condensed state. We expect that Hi-C matrices of condensed holocentric chromosomes would present a contact pattern like TADs. This proposed mechanism brings not only the centromeric nucleosomes to a linear organization, but the chromatin loops in between them as well. By regulating the distribution of LEs and centromeric nucleosomes, the same mechanism could form different structural arrangements and compaction levels. We observed that the modelled holocentric or monocentric-like distribution of centromeric nucleosomes led to different chromosome lengths (Supplementary Figure 2).

We further suggest that the presence of kinetochores in holocentric species has a direct impact on the chromosome conformation (Figure 7). With a model of the kinetochore plate bound to the holocentric line we observed a constriction to the chromatin loops, forming a groove-like structure along the chromatid. The presence of kinetochores also limited, possibly shortened, the length of the centromeric line.

Despite the small size of the chromosome that we simulated, we expect that larger holocentric chromosomes are subjected to the same mechanism, but its compaction may require different amounts of LEs or result in different chromatin loop lengths. Likewise, we expect the formation of the centromeric groove-like structure to depend on the chromatin loop lengths and the arrangement of the kinetochores.

Limitations of our model lie in the simplicity of our assumptions. Here, centromeric nucleosomes can either block or not, either anchor or not the LEs. But these effects may be milder and not always block the motion of LEs (47), or only temporarily anchor the LEs. The loop extrusion mechanism lacks the possibility of LEs traversing each other (49). And we could infer more about the effects of centromeric nucleosomes simulating other distributions and the presence of eu-and heterochromatic regions (50). But we expect that our proposed mechanism will generally apply to centromeres and its improvements will lead to the observed variety of centromeric structures. Overall, none of the mechanisms we described is novel, but their combined action according to our dynamic model could account for the structural and evolutional diversity of the different centromere types.

## DATA AVAILABILITY

All the Fortran and Python scripts, for running and analyzing the simulations, are available in bitbucket.org/ipkdg/polymer_simulations.git.

## SUPPLEMENTARY DATA

Supplementary Data are available at NAR online.

## ACKNOWLEDGEMENTS

We would like to thank the research groups Domestication Genomics and Chromosome Structure and Function at IPK for fruitful discussions and critical comments.

## FUNDING

This work was supported by the German Research Foundation [MA 6611/4-1 to A.S.C and M.M., HO 1779/31-19 to A.H. and Schu 762/11-1 to V.S.].

## CONFLICT OF INTEREST

No conflict of interest exists.

## Notes

https://doi.ipk-gatersleben.de/DOI/bfd1aba2-86e2-4b64-9825-fa795178080a/7e2741cd-5d8a-49ab-8f11-646cc6a4d36f/2

### Competing Interest Statement

The authors have declared no competing interest.

