## Supplementary Data for "A simple model explains the cell cycle-dependent assembly of centromeric nucleosomes in holocentric species"

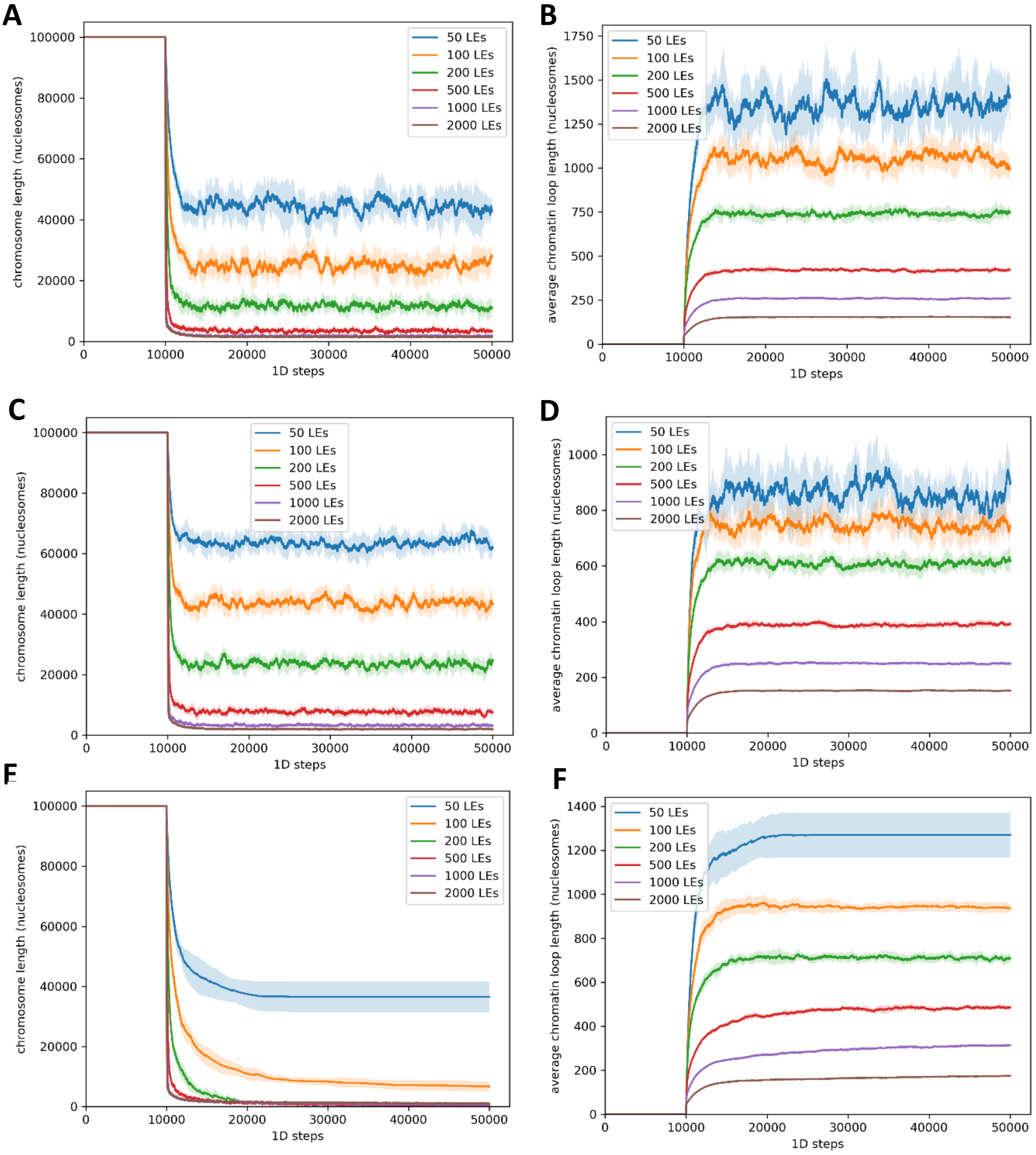


Supplementary Figure 1. Chromosome length (A, C, E) and average chromatin loop length (B, D, F) over time for different interaction effects of centromeric nucleosomes with LEs: (A, B) no effect; (C, D) blocking effect; and (E, F) anchoring effect. Simulations with different amounts of LEs are shown in different colours: 50 LEs in blue, 100 in orange, 200 in green, 500 in red, 1000 in lilac and 2000 in brown. A plateau for the final steps of the simulation means these two parameters have reached equilibrium.

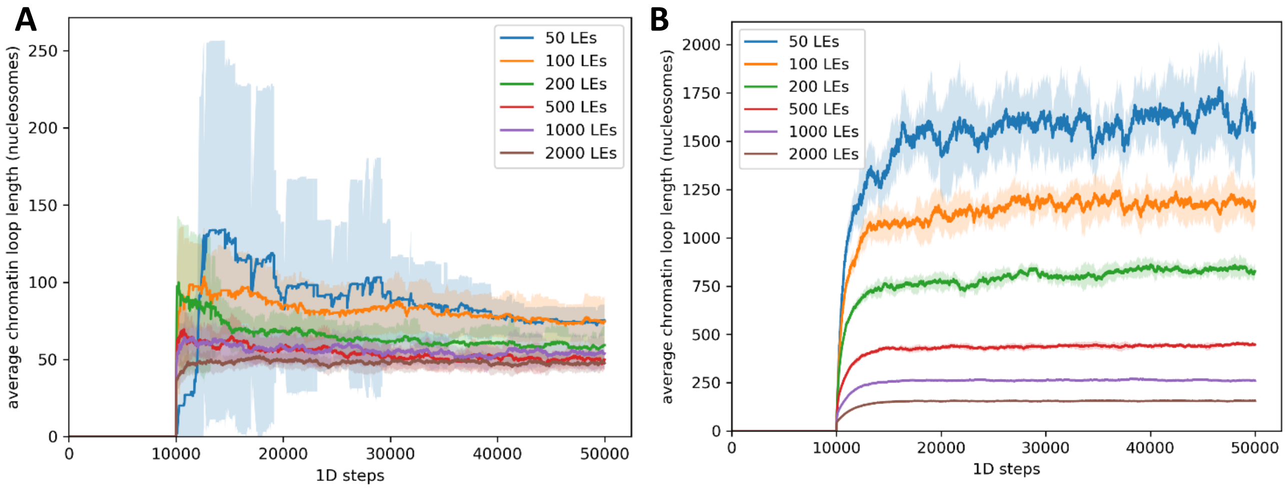


Supplementary Figure 2. Average chromatin loop length over time for (A) the centromeric region, and (B) outside the centromeric region. Simulations with different amounts of LEs are shown in different colours: 50 LEs in blue, 100 in orange, 200 in green, 500 in red, 1000 in lilac and 2000 in brown. A plateau for the final steps of the simulation means the length has reached equilibrium.

Supplementary Movie 1. Condensation of a holocentric chromosome, where the centromeric nucleosomes have no effect on the Loop Extruders. Three visualization modes are present. Chromatin is shown as a 10 nm thick grey fibre, the Loop Extruders are represented by the bound nucleosomes as pairs of yellow beads and the centromeric nucleosomes are shown as beads coloured in red or in a gradient from blue to red, following their chromosome position, as indicated by the subtitles. The simulation presents 1000 LEs and 100 centromeric nucleosomes. This movie is available as HolocentricChromosome_noEffectCentromeres.mp4 with DOI 10.5447/ipk/2021/5.

Supplementary Movie 2. Condensation of a holocentric chromosome, where the centromeric nucleosomes have a blocking effect on the Loop Extruders. Three visualization modes are present. Chromatin is shown as a 10 nm thick grey fibre, the Loop Extruders are represented by the bound nucleosomes as pairs of yellow beads and the centromeric nucleosomes are shown as beads coloured in red or in a gradient from blue to red, following their chromosome position, as indicated by the subtitles. The simulation presents 1000 LEs and 100 centromeric nucleosomes. This movie is available as HolocentricChromosome_blockingCentromeres.mp4 with DOI 10.5447/ipk/2021/5.

Supplementary Movie 3. Condensation of a holocentric chromosome, where the centromeric nucleosomes have an anchoring effect on the Loop Extruders. Three visualization modes are present. Chromatin is shown as a 10 nm thick grey fibre, the Loop Extruders are represented by the bound nucleosomes as pairs of yellow beads and the centromeric nucleosomes are shown as beads coloured in red or in a gradient from blue to red, following their chromosome position, as indicated by the subtitles. The simulation presents 1000 LEs and 100 centromeric nucleosomes. This movie is available as HolocentricChromosome_anchoringCentromeres.mp4 with DOI 10.5447/ipk/2021/5.

Supplementary Movie 4. Condensation of a monocentric chromosome, where the centromeric nucleosomes have an anchoring effect on the Loop Extruders. Three visualization modes are present. Chromatin is shown as a 10 nm thick grey fibre, the Loop Extruders are represented by the bound nucleosomes as pairs of yellow beads and the centromeric nucleosomes are shown as beads coloured in red or in a gradient from blue to red, following their chromosome position, as indicated by the subtitles. The simulation presents 1000 LEs and 20 centromeric nucleosomes. This movie is available as MonocentricChromosome_anchoringCentromeres.mp4 with DOI 10.5447/ipk/2021/5.
